# Ultra-compact Dual-band Smart NEMS Magnetoelectric Antennas for Simultaneous Wireless Energy Harvesting and Magnetic Field Sensing

**DOI:** 10.1101/2020.06.22.165894

**Authors:** Mohsen Zaeimbashi, Mehdi Nasrollahpour, Adam Khalifa, Anthony Romano, Xianfeng Liang, Huaihao Chen, Neville Sun, Alexei Matyushov, Hwaider Lin, Cunzheng Dong, Ziyue Xu, Ankit Mittal, Isabel Martos-Repath, Gaurav Jha, Nikita Mirchandani, Diptashree Das, Marvin Onabajo, Aatmesh Shrivastava, Sydney Cash, Nian X. Sun

## Abstract

Ultra-compact wireless implantable medical devices (IMDs) are in great demand for healthcare applications, in particular for neural recording and stimulation. Current implantable technologies based on miniaturized micro-coils suffer from low wireless power transfer efficiency (PTE) and are not always compliant with the specific absorption rate imposed by the Federal Communications Commission, particularly for deep brain implantation where field attenuation and tissue loss are significant. Moreover, current implantable devices are reliant on recordings of voltage or current. This has two major weaknesses: 1) the necessary direct contact between electrode and tissue degrades over time due to electrochemical fouling and tissue reactions, and 2) the necessity for differential recordings across space. Here, we report, for the first time, an ultra-compact dual-band smart nanoelectromechanical systems magnetoelectric (ME) antenna with a size of 250×174 μm^2^ that can efficiently perform wireless energy harvesting and sense ultra-small magnetic fields such as those arising from neural activities. The proposed smart ME antenna has a wireless PTE 1~2 orders of magnitude higher than any other reported miniaturized micro-coil, allowing the wireless IMDs to be compliant with the specific absorption rate (SAR) limit and to operate under safe exposure of radio frequency energy. Furthermore, the magnetic sensing capability of the proposed smart ME antenna, with a limit of detection of 300~500pT at > 200Hz, should allow the IMDs to record neural magnetic fields from the brain without requiring differential recording.

## 1- Introduction

Direct measurement of widespread neuronal activity at the level of small ensembles of neurons is essential for both basic research and clinical applications. Current technology for doing so exists, but is limited in the number of sites and spatial extent coverable. To address the need for a distributed neural interfacing tool, a new generation of wirelessly powered standalone implantable medical devices (IMDs) have emerged in the last few years [1–4]. Wirelessly powered IMDs eliminate the invasiveness and discomfort caused by batteries and wires in most conventional implants. The two major wireless powering modalities today are electromagnetic and ultrasonic. Regardless of the powering method used, as the receiver shrinks, it becomes increasingly difficult to deliver sufficient power to operate the IMD, which can range from a few to several hundred microwatts depending on the application. This puts a heavy burden on the miniaturization of coils and piezo devices, which explains why many state-of-the-art single channel devices remain bulky [5–9]. Ultimately, these devices will need to be surgically implanted into the brain but the technology remains too large for human applications. Some untethered IMDs have successfully miniaturized their transducers [10–14] down to the scale of a human hair diameter. However, the current prototypes do not have the necessary wireless efficiency to be safely powered in human applications because the SAR limit imposed by the FCC needs to be respected. In this article, a novel acoustically actuated nanomechanical ME antenna with the highest power transfer efficiency reported to date is described. These ME antennas incorporate a magnetic and piezoelectric heterostructure where the magnetic film senses H-components of EM waves. The magnetic layer then induces an oscillating strain, which generates a piezoelectric voltage output at the electromechanical resonance frequency. By exploiting this transduction mechanism, ME antennas do not suffer from the same miniaturization constraints as coils, and they can be driven by weak magnetic fields [15].

We have most recently reported magnetoelectric (ME) nanoelectromechanical system (NEMS) resonators as single-band ME antennas for wireless communication [15], and ME sensors for quasi-static low-frequency magnetic field sensing [16] in two separate devices with two different designs and different operating frequencies. The ME antenna is based on ME FBAR (thin film bulk acoustic wave resonator) working at 2.5 GHz; while the ME sensor is based on a ME NPR (nano-plate resonator) with interdigitated electrode and an operation frequency of 215MHz. In this paper we present the first ever smart ME antenna with unprecedented characteristics that are ideal for IMDs: (1) ultra-compact antenna for highly efficient wireless power transfer efficiency and data communication at GHz; (2) ultra-sensitive magnetometer capable of sensing picoTesla low-frequency fields by using MHz resonance; and (3) simultaneous operation at two different frequency bands, GHz for wireless power transfer and data communication, and MHz for magnetic field sensing. Compared to neural electrical sensing based on differential voltages from neural probes or wireless implants [17], the proposed smart ME antennas operate based on neuronal magnetic sensing, providing multiple advantages: (1) magnetic neural sensing is not differential, which enables more compact neural recording implants or elements [17]; (2) thanks to its physical characteristics, magnetic neural sensing is more localized than electrical sensing, which consequently enables better spatial resolution of neuronal activities; (3) it is practical to create safe and contact-less implants coated with bio-compatible polymer films such as parylene; (4) the same technology can be used for animals and human, allowing for direct comparisons and easier translation of animal to human information; and (5) compact size allows for distributed, addressable, high channel count use in the parenchyma, pial surface, extra dural, and in both central and peripheral nervous tissue [18].

## 2- Mechanically Actuated Magnetoelectric Antennas

Magnetoelectric antennas are based on multiferroic materials, which are materials that show more than one of the primary ferroics properties. These primary ferroics include: a) ferromagnetism, a phenomenon in which magnetization can be changed by an applied magnetic field; b) ferroelectricity, in which electrical polarization can be altered by an applied electric field; c) ferroelasticity, in which a deformation can be changed by an applied stress. ME antennas, consisting both piezoelectric and magnetostrictive thin-film materials, exploit the aforementioned ferroic properties in order to receive and transmit electromagnetic waves. Piezoelectric thin-film couples electrical polarization and mechanical strain, and magnetostrictive material couples magnetic polarization and mechanical strain. A piezoelectric and magnetostrictive heterostructure, deposited by sputtering system is used to couple the discussed ferroic orders. The diagram in Fig. 1a visualizes the concepts of the transmitter (T_X_) and receiver (Rx) operating modes of an ME antenna, where the top and bottom rectangular boxes represent the mechanically-coupled magnetostrctive and piezoelectric layers, respectively. In T_X_ mode (L), an RF voltage is applied to piezo material, which in turn generates mechanical strain. This mechanical strain is then transferred to the magnetostrictive thin-film, which subsequently radiates EM wave due to a piezomagnetic phenomenon. In Rx mode (R), the same sequence occurs in reversed order, where incoming EM wave towards magnetic material generates strain. The strain is then transferred to the piezoelectric thin-film, which in turn generates an RF voltage in the output. The simulation results of strain and displacement distribution in magnetostrctive and piezoelectric layers are shown in Appendix A. The incoming EM wave produces a strain in magnetostrctive layer; the strain is then transferred to piezoelectric layer, which in turn induces a voltage across its thickness that can be used for energy harvesting purposes.

**Figure 1.**
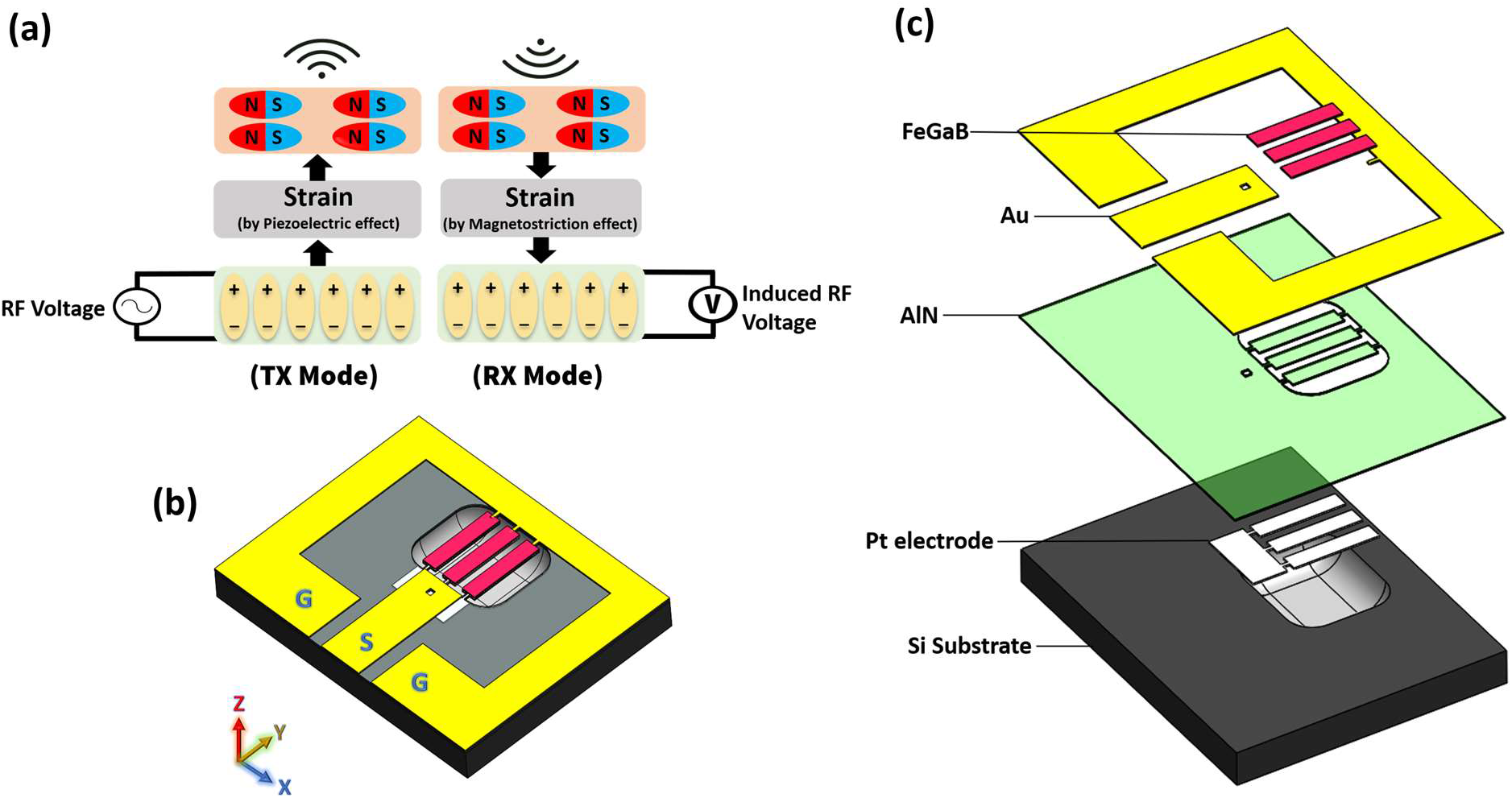
a) Diagram showing the concept of Tx and Rx operating modes of ME antenna, where the top and bottom rectangular boxes represent the magnetostrctive and piezoelectric layers, respectively; (b) 3D schematic of the ME antenna on released Si substrate. The yellow color indicates a gold ground ring and GSG pads that are used for probing and wire-bonding the antenna to PCB; (c) exploded structure of the ME antenna showing all the layers.

Fig. 1b and 1c show the 3D schematic of the ME antenna from this work, for which fabrication steps are summarized in Appendix B. This ME device consists of three parallel rectangular ME resonators, each with a size of 250×50 μm^2^, giving an overall size of 250×174 μm^2^, including two 12 μm gaps between resonators. The Si substrate underneath the ME resonators is etched in order to release the ME elements and prevent damping of the acoustic vibration, a process that improves the performance of ME antenna. The resonance frequency of the rectangular ME elements is defined by the size of piezoelectric thin film materials. AlN was used as the piezoelectric phase and FeGaB as magnetostrictive phase that are mechanically coupled through the interface. A rectangular thin-film piezoelectric material can exhibit three resonance frequencies: along width, length, and thickness. Here, we use the width and thickness modes of the ME resonators since they are more efficient than the length mode. Therefore, the proposed ME antenna has two operational resonance frequencies: (1) a nano-plate resonator (NPR) frequency corresponding to the contour (width) mode of vibration in piezoelectric thin-film, and can be expressed as 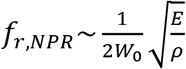, where W_0_ is the width of the resonator, and E and ρ are the equivalent Young’s modulus and equivalent density of the AlN, respectively; (2) a thin-film bulk acoustic wave resonator (FBAR) frequency associated with the thickness mode of vibration applied to the resonator, which can be expressed as 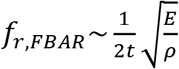, where t is the thickness of the AlN thin-film. Since the designed antenna operates in two different frequency bands, we refer to this device a smart or hybrid FBAR/NPR ME antenna. It is notable that since the thickness of the AlN film is much smaller than its width, the thickness mode (FBAR) resonance frequency of the ME antenna is significantly higher than the width mode (NPR) resonance frequency.

Fig. 2a shows an optical microscope image of a fabricated smart ME antenna with three parallel ME resonators connected to a gold ground ring with GSG probing pads. Three parallel ME resonators were used for enhanced wireless energy transfer efficiency without significantly enhanced device size. The red color is due to the reflection from the AlN thin-film because the Si substrate underneath is etched. Fig. 2b displays the measured S11 of the thickness mode of the smart ME antenna showing a resonance frequency at 2.51 GHz, which corresponds to 500 nm thickness of AlN. The two peaks are slightly apart from each other as a result of the small difference in the stress levels of the three rectangular ME resonator array. Fig. 2c shows the S11 of the width mode showing a resonance frequency at 63.6 MHz, which corresponds to 50 μm width of AlN. It is noteworthy that the S11 plots of both thickness and width modes were measured after wire bonding the ME antenna to a 200 μm thick paper PCB, on which an SMA connector is mounted in order to connect the ME antenna to a vector network analyzer (VNA). We observed some impedance matching (S11) degradation after the wire bonding process compared to the case of directly probing ME antenna pads using standard GSG probes. This change is attributed to the extra series inductance and resistance added by bonding wires. The insets in Fig. 2b and 2c show the equivalent Modified Butterworth–Van Dyke (MBVD) circuit model for thickness and width modes of ME antenna, respectively. In the following sections it will be shown that thickness and width modes of smart ME antenna exhibit a high performance while harvesting the RF energy and sensing magnetic fields, respectively—functionalities that can be utilized in ultra-miniaturized brain and body implantable devices and wearable technologies.

**Figure 2.**
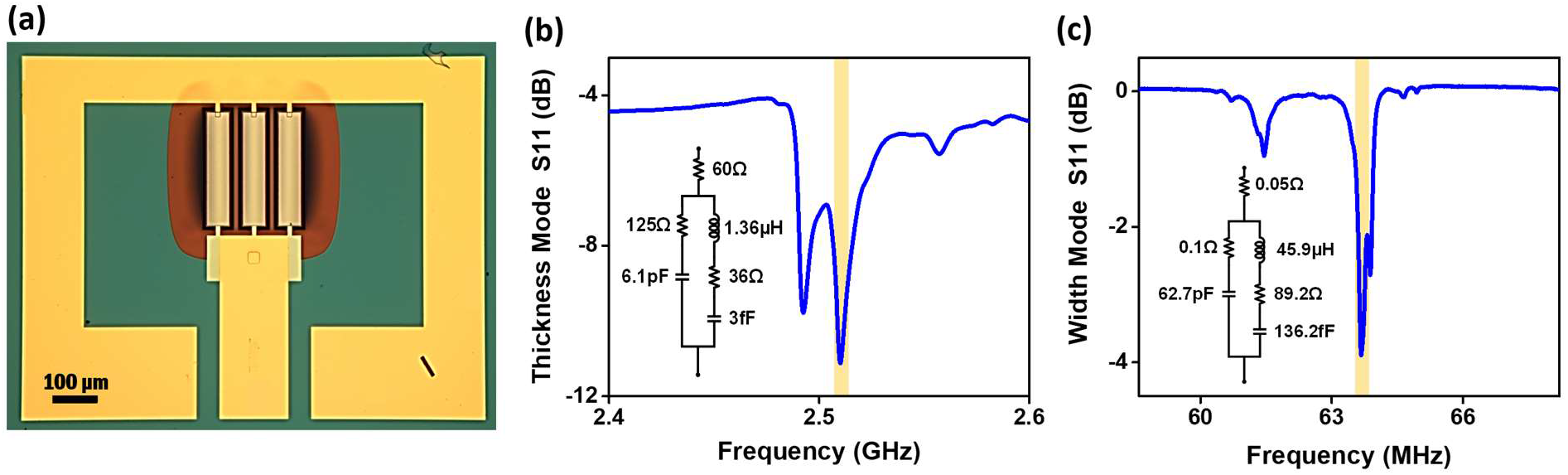
(a) Optical microscope image of a fabricated smart ME antenna; (b) S11 of the thickness mode of the smart ME antenna showing a resonance frequency at 2.51GHz; (c) S11 of the width mode of the smart ME antenna showing a resonance frequency at 63.6MHz. The insets in (b) and (c) show the equivalent MBVD circuit model of ME antenna in each mode.

### 2-1- Wireless Energy Harvesting with Smart ME Antennas

An ME antenna’s energy harvesting mode functions based on the Rx mode diagram in Fig. 1a, where an incoming radio frequency magnetic field induces strain in the magnetostrcitive material (FeGaB). This strain is then transferred to the piezoelectric thin-film, and finally the piezoelectric material generates an RF voltage at the output that can be used to power electronic circuitry of the medical implantable device. As mentioned before, the thickness mode of the ME antenna with a resonance frequency of 2.51 GHz is used for energy harvesting because it outperforms the width mode for this application, while the width mode of the smart ME antenna exhibits higher magnetic field sensitivity.

We have designed a single turn T_X_ coil on an FR4 PCB to investigate the energy harvesting performance and efficiency of ME antennas. The Sonnet simulation toolbox was used to optimize the Tx coil in terms of size, Q-factor, self-resonance frequency (SRF), and trace width. The optimized and fabricated Tx coil, shown in Fig. 3a, has a self-resonance frequency (SRF) of 4.1 GHz, Q-factor and inductance of 110 and 10.49 nH (both @ 2.51 GHz and in air), length of 10 mm, and trace width of 3 mm. An L-match capacitive network, also displayed in Fig. 3a, is used to match the Tx coil at 2.51 GHz for maximization of the transmitted power. An SMP connector is soldered to the PCB to connect the Tx coil to the VNA for impedance matching and characterization. Fig. 3b shows the reflection coefficient (S_11_) of the T_X_ coil matched at 2.51GHz, which is the FBAR resonance frequency of the smart ME antenna as seen in Fig. 2b. Note that S11 in Fig. 3b was measured when the Tx coil was surrounded by air. When the Tx coil is placed close to the skin or tissue, the resonance frequency of the matched Tx coil would shift by 100-300 MHz compared to the air medium; therefore, one has to re-match the Tx coil when tissue is present. For this reason, the matching network in Fig. 3a has been implemented with variable capacitors. As mentioned earlier, the GSG pads of a smart FBAR/NPR ME antenna, shown in Fig. 1b, are wire-bonded to a PCB, and an SMA connector is mounted on the PCB to obtain access to ME antenna’s pads.

**Figure 3.**
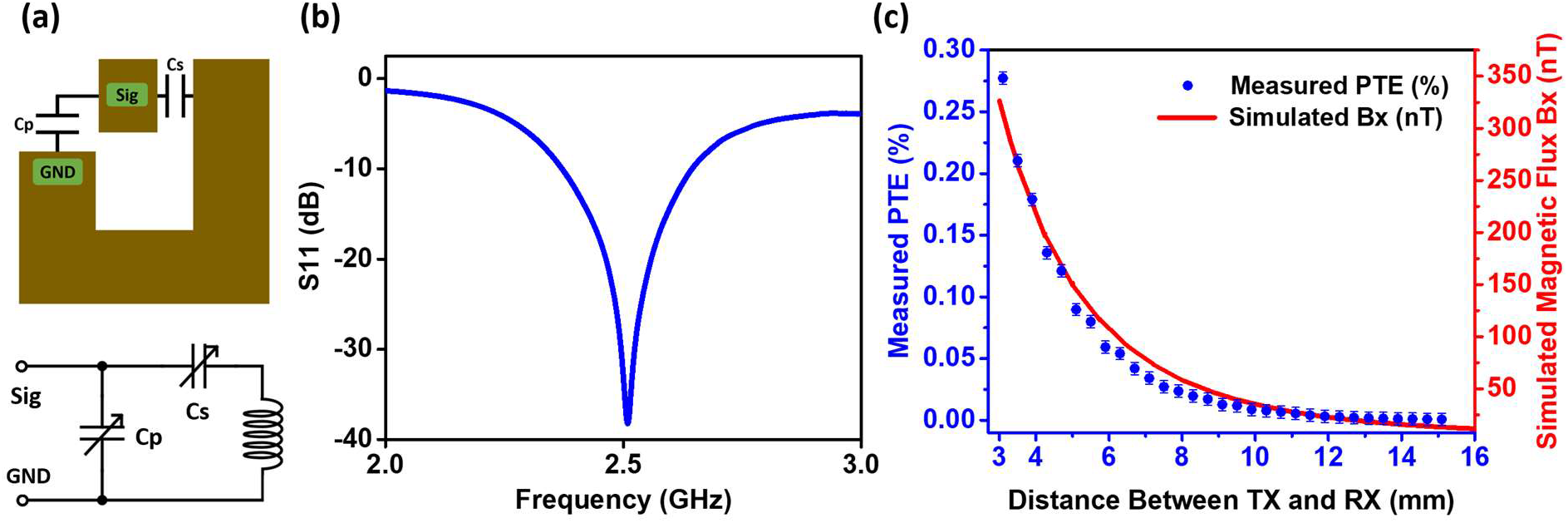
(a) Diagram of the PCB traces of the Tx coil used in the energy harvesting experiment, and schematic of the L-matching network for impedance matching of the coil; (b) measured S11 of the Tx coil matched at 2.51 GHz, which is the FBAR resonance frequency of the smart ME antenna. (c) The left-axis shows the measured PTE versus distance between ME antenna and Tx coil, and the right axis shows the simulated magnetic flux density along the x-axis (Bx) generated by the Tx coil versus distance from Tx coil.

The diagram of the experimental setup for the energy harvesting measurements is shown in Appendix E, Fig. S5. The ME antenna and Tx coil are placed on two 3D printed plastic manipulators in order to precisely control their position. Plastic manipulators, instead of metallic ones, are used to minimize the effect of objects surrounding the Tx coil and their impact on S11 of the coil. The manipulators are adjustable along the X, Y, and Z axes, and have a spatial resolution of less than 200 μm, allowing an accurate investigation of impacts of distance and misalignment between the ME antenna and Tx coil. To measure the power transfer efficiency (i.e., the ratio of received power by the ME antenna to transmitted power from the Tx coil), the ME antenna was connected to a power spectrum analyzer, and the Tx coil was connected directly to a VNA under 7 dBm power. The left-axis in Fig. 3c shows the measured PTE versus distance between ME antenna and Tx coil, and right axis shows the simulated magnetic flux density along the x-axis (Bx) generated by the Tx coil versus distance from the Tx coil. The Cartesian coordinate system here is the same as that in Fig. 1b. It is important to note that the ME antenna is sensitive along the x-axis due to the direction of the FeGaB material’s easy axis, which is defined based on the applied DC field during the deposition process of this layer. Therefore, the ME antenna only responds to the magnetic fields along the x-axis when generating a voltage. Fig. 3c shows that the rate of reduction of the measured PTE and the simulated magnetic flux density along the x-axis (Bx) are matching, and both are decreasing approximately cubically with the distance. The Tx coil was simulated with the COMSOL AC/DC module at 2.51 GHz under an input current of 10 mA, which corresponds to 7 dBm power (with 50 Ohm resonance condition as in this experiment).

### 2-2- Misalignment and Rotation of the Smart ME Antenna During Energy Harvesting

Fig. 4a shows the simulated PTE of a ME antenna in the XY plane at a fixed distance of z = 8 mm from Tx coil. In other words, this plot shows PTE versus misalignment of the ME antenna with respect to Tx coil. For this plot, the PTE was measured at 361 points (array of 19×19 pixels) with a step size of 1.6 mm along both X and Y axis, giving a total measurement range of 14.4× 14.4 mm^2^. The black outline inside the plot identifies the position of the Tx coil within the XY plane. The measurement results show that the PTE, which is proportional to the amplitude of the magnetic flux density (Bx) to which the ME antenna is sensitive, reaches a maximum above the PCB traces along the Y axis, which produces a magnetic flux density along the X axis. Furthermore, the magnetic flux focal size above the longer PCB trace is much larger than the focal point size above the shorter PCB trace. It is noteworthy that the ME antenna was in a fixed position during this measurement, and the Tx coil was shifted along the X and Y axes, which, due to reciprocity theory of electromagnetics, is equivalent to fixing the Tx coil and moving the ME antenna along X and Y axis. Fig. 4b shows the COMSOL simulation results for the magnetic flux (Bx) in the XY plane at z = 8mm distance from the coil. The black outline shows the position of the Tx coil within the XY plane. The simulation results confirm the PTE measurements, and they show that the longer PCB trace on the right side of the Tx coil produces an 11× larger magnetic flux focal point than the shorter trace on the left, while this trace is only about 2× longer than the shorter trace.

**Figure 4.**
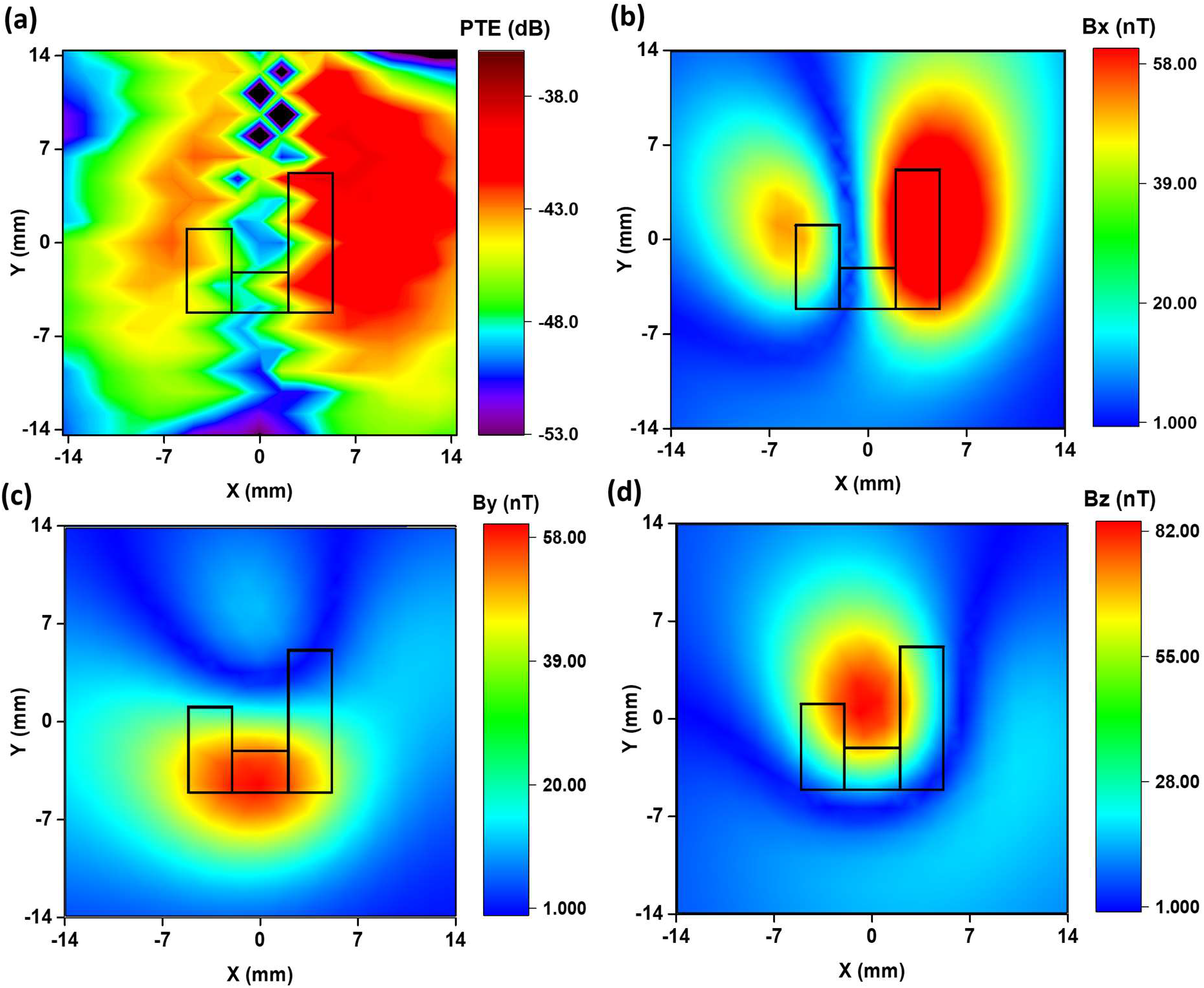
(a) Measured PTE of the ME antenna in the XY plane at a fixed distance of z = 8 mm from the Tx coil. The black figure inside the plot shows the coordinate of the Tx coil in the XY plane; (b) COMSOL simulation results of magnetic flux (Bx) in the XY plane at z = 8mm distance from the coil. The simulation results confirm the PTE measurements and show that the longer PCB trace on the right side of Tx coil produces a much larger (> 11×) magnetic flux focal point than the shorter trace on the left; (c) and (d) show the magnetic flux densities By and Bz in the XY plane at z = 8 mm distance from the Tx coil, respectively. The magnetic flux By reaches a maximum above the lower copper trace because the current is flowing along the X axis in this direction. The magnetic flux Bz is peaking at the center of the Tx coil because the field loops from copper traces are constructively interfering and giving a field stronger than Bx and By.

In biomedical applications, when an implantable device is inserted into the body or brain, inevitably that device will move and rotate after its initial placement. In such a situation with a change of orientation, and considering that the ME antenna is only sensitive to magnetic flux fields along the X axis, it is crucial to ensure that the external Tx coil will be able to power the implanted device even if it is oriented towards the Y or Z axes, or any other direction in-between. Simulation results show that the designed Tx coil, or any other rectangular shaped coil, is able to produce magnetic flux density along almost every direction, including X, Y, Z, XY, etc. This is because the rectangular Tx coil has trace lines along both X and Y axes, each producing magnetic flux along the Y and X directions, respectively. Moreover, since the Tx coil is a one-turn loop, the magnetic fluxes generated by copper traces constructively interfere at the center of the coil, and produce an even stronger magnetic field along the Z axis. Fig. 4c and 4d show the magnetic flux densities By and Bz in the XY plane at z = 8mm distance from Tx coil. The magnetic flux By reaches a maximum above the lower copper trace because the current is flowing along X the axis in this direction. The magnetic flux Bz is peaking at the center of the Tx coil because the field loops from copper traces are constructively interfering and giving a field stronger than Bx and By. The magnetic flux distribution along several other directions is available in Fig. S3 in Appendix C. Fig. S3b, for instance, shows the magnetic field along the y = x line — i.e. 45° away from both x and y axis — in the XY plane at z = 8mm distance from T_X_ coil. The fields along both X and Y axis, each with a 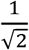 factor contribution, produce the magnetic fields along this line.

Based on the discussed results, we can safely conclude that, regardless of how the ME antenna moves and its orientation after implanting the device, by scanning the T_X_ coil one can align the peak of magnetic flux to the ME antenna and power the implant. For instance, if the ME antenna is rotated in a way that its sensitive direction is along the Z axis, the center of the T_X_ coil should be aligned to the implant to power the device.

### 2-3- In-vitro Wireless Energy Harvesting Experiment on Mice Brain Tissue

The discussion in the previous section was on the wireless energy harvesting performance of ME antenna in the air medium. In real biomedical applications, however, it important to take into account the effect of the tissue medium because tissue, when placed between Tx and Rx antennas, can impact the impedance matching of Tx coil and also the magnetic field distribution. In order to investigate the tissue impact on wireless harvesting tests we have completed experiments in which mice tissue was placed between the Tx coil and the ME antenna. The mice tissue was 6 mm thick, as shown in Fig. 5a, and included all the layers including scalp, skull, meninges layers (dura, arachnoid, pia mater), and grey mater. The tissue was placed on top of the Tx coil with 1.6 mm air gap in between, and the ME antenna was precisely aligned on top of the tissue, with 1 mm air gap, to get the maximum power. The total thickness between Tx coil and ME antenna was 8.6 mm. It is notable that when the tissue was placed on top of the Tx coil, the resonance frequency of Tx coil was shifted downward by about 100-300 MHz, which is attributed to the changes of the boundary conditions and also of the impedance seen by Tx coil compared to the air medium. Therefore, the Tx coil was re-matched using an L-match capacitive network after placing the tissue to ensure maximum power is transmitted by the coil.

**Figure 5.**
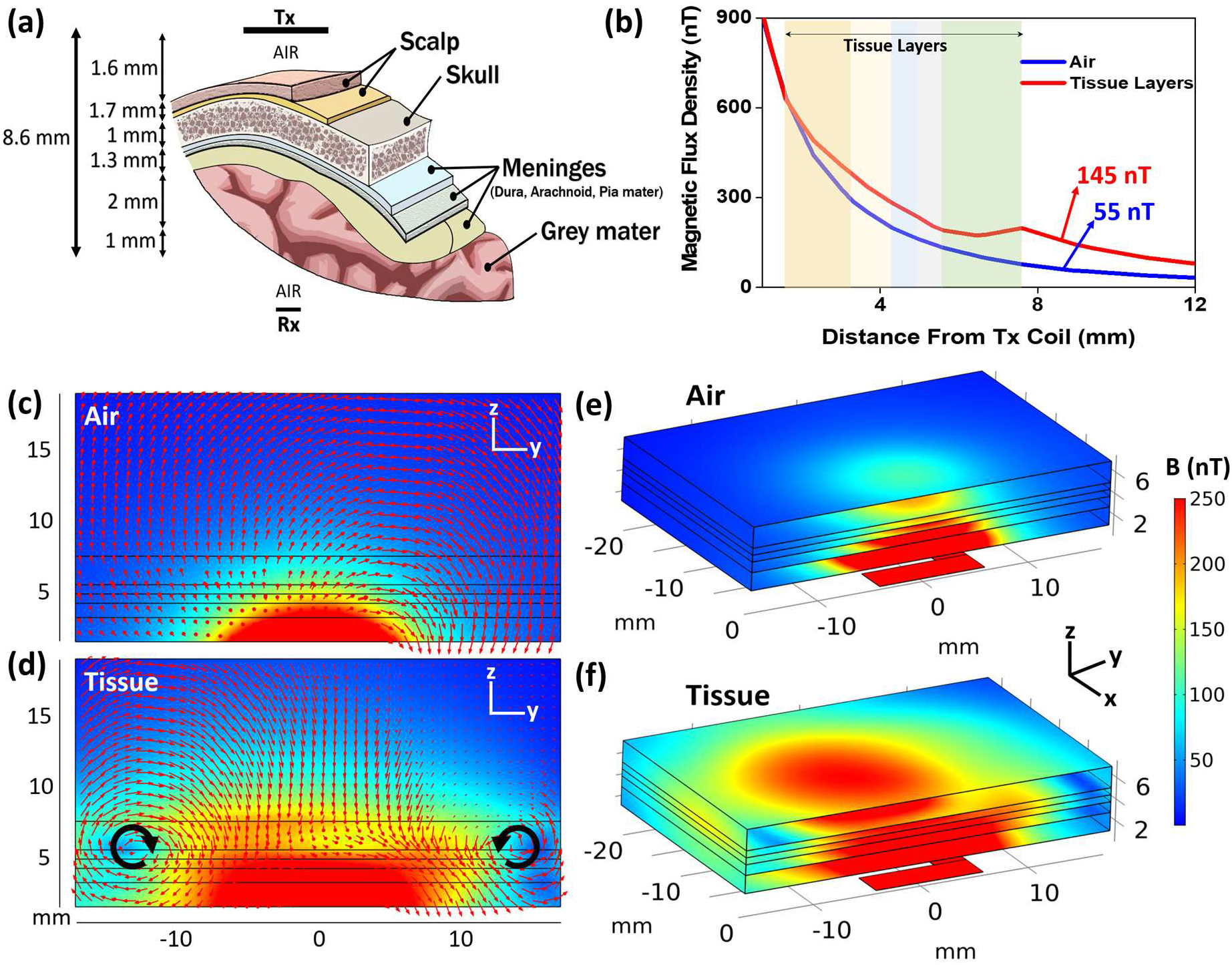
(a) Schematic of mice tissue layers used in in-vitro energy harvesting experiment; (b) Magnetic flux density vs distance from Tx coil for tissue layers and air medium, showing that the magnetic flux density is stronger when tissue is present. Color bands show the coordinate of different tissue layers; (c) and (d) simulation results showing the magnetic field distribution and vectors on a cross-section cut of air and tissue mediums, respectively. The black lines show the coordinate of tissue layers. The two vortexes in 6d are due to the magnetic fields generated by the eddy current loops inside the meninges layer, where tissue conductivity is maximum. Tx coil is located at the bottom. The y-axis shows the distance from Tx coil; (e) and (f) show the perspective view of magnetic field distribution on a cross-section cut inside air and tissue mediums, respectively. The results in (c), (d), (e), and (f) show that the magnetic field distribution in the space changes and is higher when a tissue is present. The color bar scale on the right is valid for all figures.

It turns out that the measured wireless PTE of the system is about 3.1 times higher when the tissue is present compared to air medium, which was against our initial intuition because the tissue could add some loss to the system and in fact reduce the PTE. In order to shed light on this observation, we simulated and compared the tissue and air mediums using the COMSOL AC/DC module under the same current and frequency conditions as our experiment. Simulation results also matched the experiments and showed that the magnetic flux density at 8.6 mm distance is higher when the tissue is present. Fig. 5b shows the magnetic flux density versus the distance from the Tx coil for the air and mice tissue models. The simulation data shows that the magnetic flux density at 8.6 mm is 2.6 times higher when tissue is present compared to air medium. This field enhancement is due to the fact that EM wave is in propagation mode inside the tissue rather than in evanescent near field coupling mode where magnetic field reduces more rapidly. Since the relative permittivity of the tissue is 40-60 times higher than that of air at 2.51 GHz, the EM wave travels slower in the tissue medium and therefore its effective wavelength is shorter. Hence, an EM wave can be propagative even in regions very close to the coil. It is notable that Ho et al. in [19] have observed the same phenomena where the EM field is stronger when a tissue medium is present. Fig. 5c and 5d show the magnetic flux distribution and vectors on a cross-section of air and tissue layers, respectively. The black lines show the coordinate of the tissue layers. The magnetic field distribution and vector flows are normal in the air medium, but they are distorted and noticeably changed in the tissue medium. As it is shown, there are two vortexes in the tissue medium at the interface of the meninges layer, where conductivity of the tissue is the highest. The reason for these vortexes is that the Bz magnetic field created by Tx coil generates eddy current loops inside the tissue layers, particularly in the meninges layers because of their high conductivity, and these eddy currents loops which exist on x-y plane create their own magnetic flux densities. The magnetic fields created by eddy current loops inside the tissue interfere with the Tx coil magnetic fields, creating the vortexes and distorting the magnetic field distribution compared to the air medium. More detailed explanation on the generated eddy currents in the tissue is available in Appendix D. Fig. 5e and 5f show the 3D view of magnetic field distribution in air and tissue mediums, respectively. It can be observed that the magnetic flux density is higher in the tissue medium, which matches the experimental observation.

Table 1 compares the ME antenna’s energy harvesting performance with that of five other state-of-the-art works selected because of their sub-mm size. These other systems, to the best of our knowledge, are the best published small-size micro-coils. The table compares different systems in terms of PTE, size of energy harvesting element (micro-coil or ME antenna), and the distance between Rx and Tx, as well as the overall energy harvesting performance using the figure of merit (FOM) from [20] that is defined as follows:

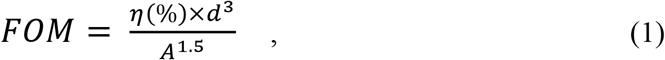

where *η* is the PTE as percentage, *d* is the distance between Rx and Tx in mm, and *A* is the area of the Rx element (coil or ME antenna) in *mm*^2^. As it can be observed in Table 1, the FOM of the ME antenna is better than that of other mm-scale coils (row 1 in table) and significantly superior to other μm-scale coils (row 2-5 in the table).

**Table 1.** Comparison of the ME antenna’s energy harvesting performance with that of five other published micro-coils with sub-mm size. The formula in equation (1) is used to calculate the figure of merit. This formula considers the area of the receiver, the distance between Rx and Tx, and the wireless power transfer efficiency.

| Ref. | Rx Area<br>(mm<sup>2</sup>) | Rx Type | Tx-Rx<br>Distance<br>(mm) | Wireless<br>System | Medium | PTE (%)<br>= $\eta_{Tx}$<br>* *Path Loss*<br>* $\eta_{Rx}$ | FOM |
| --- | --- | --- | --- | --- | --- | --- | --- |
| Dukju [25] | 0.79 | Off-chip coil | 12 | 2-coil | Tissue (beef) | 0.56 [200MHz,<br>PDL=224μW] | 1378 |
| Khalifa [11] | 0.09 | On-chip coil | 6.6 | 2-coil | Tissue (lamb) | 0.01* | 106 |
| Liuqing [26] | 0.01 | On-chip coil | 0.5 | 3-coil | Air | 0.12 [2GHz] | 12 |
| Nai-Chung Kuo<br>[27] | 0.01 | On-chip coil | 2.2 | 2-coil | Air | 0.0079* [4.8GHz,<br>PDL=100μW] | 84 |
| ***This work***<br>***(Smart ME***<br>***antenna)*** | **0.044** | **ME antenna** | **8.6** | **1-coil + ME**<br>**antenna** | **Mice Head**<br>**Layers** | **0.054 [2.51GHz]** | **3721** |

\*Simulation

## 3- Magnetic Field Sensing with a Smart ME Antenna

Ultracompact magnetic sensors are of use in various applications such as biomedical imaging, unmanned autonomous systems, and geophysical studies. For biomedical applications in particular, such as recording magnetic fields of brain neural activities or mapping magnetic patterns of the heartbeat, an ideal magnetometer should have a high spatial and temporal resolution to reduce invasiveness and to be able to record fast-changing magnetic signals. Moreover, the magnetometer should operate well in an un-shielded and room-temperature environment. The vast majority of existing magnetometers lack at least one of the mentioned criteria, and cannot be used for biomedical applications. In this section we will discuss the underlying concept of magnetic field sensing using the proposed smart ME antenna, and will also assess the performance of this device for sensing sub-nT magnetic fields.

The magnetic sensing functionality of an ME antenna is based on a non-linear ME modulation technique [21], where the external magnetic field is modulated on the RF carrier signal that is applied to the ME antenna. As mentioned before, the in-plane mode of a smart ME antenna is used for magnetic field sensing because, based on our observations, this mode shows a significantly better performance for this purpose than the thickness mode. The three parallel ME elements in the smart ME antenna each consist of a FeGaB and AlN heterostructure, where the former acts as magnetostrictive layer and the latter acts as piezoelectric layer. The coupling between magnetic, electric, elastic, and thermal parameters of this FeGaB/AlN heterostructure can be expressed using a thermodynamic approach [22]. Assuming that the temperature is constant in our experiment, and since the external magnetic and electric fields as well as the external elastic force are all independent variables, Gibbs free energy *G*(*T*, *σ*, *H*, *E*) for multiferroic materials can be used as thermodynamic potential [22] and, accordingly, the total strain (*x*) of the system can be expressed as below:

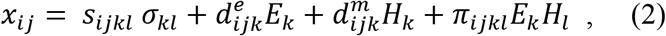

where *s_ijkl_* is the elastic compliance tensor, *σ_kl_* is the mechanical external applied stress tensor, 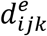 is the piezoelectric coefficient tensor of AlN thin-film, *E* is the external electric field, 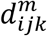 is the piezomagnetic coefficient tensor of FeGaB layer, *H* is the external applied magnetic field, and *π_ijkl_* is the piezo-coupling constant tensor. There is no external applied stress in our measurements, and therefore the first term, *s_ijkl_ σ_kl_*, which is the strain due to external mechanical stress, is almost zero and can be neglected. Thus, equation (2) can be written as below:

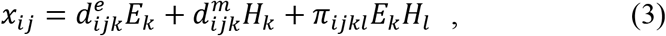

where the first term is the piezoelectric strain induced by the external electric field (RF voltage) applied to AlN thin-film, the second term is the piezomagnetic strain due to external magnetic field (the field that is being recorded), and the third term, which is the ME modulation term, is the piezo-coupling strain due to the mutual piezoelectric/piezomagnetic effects or, in other word, the strain-mediated ME effect in the AlN/FeGaB heterostructure. The external applied electric field can be written as 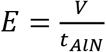, where *V* = *V_c_* cos(*ω_c_t*) and *t_AlN_* are the applied RF voltage to and thickness of the AlN layer, respectively; and the external magnetic field can be expressed as *H* = *H_m_* cos(*ω_m_t*). It is notable that *V_c_* and *ω_c_* are the amplitude and frequency of the carrier electric signal applied to ME antenna respectively, and *ω_c_* is equal to the resonance frequency of the NPR mode; *H_m_* and *ω_m_* are the amplitude and frequency of external the alternating magnetic field. The third term in equation (3), which is the strain due to ME effect, can be written as:

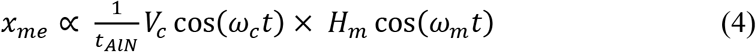

The above equation expresses the modulation between carrier voltage with frequency of *ω_c_* and modulation field with frequency of *ω_m_*. The modulation frequencies arising from the strain-mediated ME effect expressed in this equation are *ω_c_ – ω_m_* and *ω_c_* + *ω_m_*, and the amplitude of these modulation components is linearly proportional to the amplitude of external magnetic field *H_m_*, which implies the usefulness as a sensor. The linearity of the modulation components will be discussed later in this section based on Fig. 6c. Furthermore, since the low-frequency external alternating magnetic field is modulated on a high-frequency signal (at the NPR resonance frequency) and therefore its information is transferred to a higher frequency bands, the modulated component has more immunity from environmental electromagnetic, thermal, and mechanical noises that are typically all more severe in lower frequency ranges where they can easily degrade the signal-to-noise ratio (SNR) of the sensor. For instance, the work in [21] has demonstrated that a magnetoelectric modulation technique can improve the SNR of the sensor up to 72 times compared to traditional DC-biased and unmodulated sensors. It is notable that even though the mentioned work was completed on a very large magnetoelectric heterostructure, 28 mm in length, the concept and the modulation technique are similar to our scenario.

**Figure 6.**
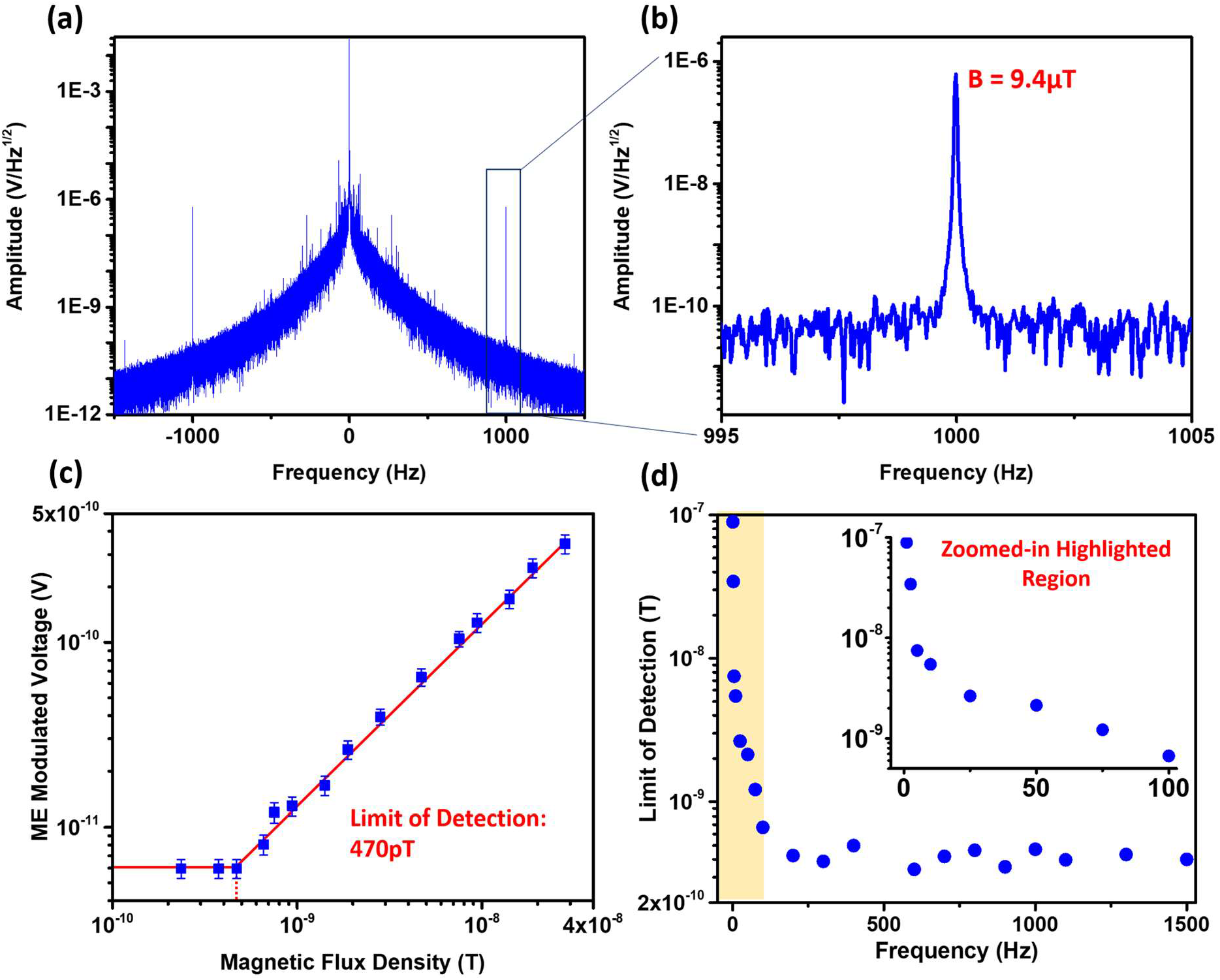
(a) Power spectrum of the reflected signal from the ME antenna after demodulation. The plot shows both modulation components of *ω_c_* – *ω_m_* and *ω_c_* + *ω_m_*. The carrier signal is located at the center and has an amplitude of 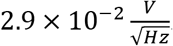. The 1/*f* electronics noise decreases significantly with increasing frequency, which improves the SNR of the sensor at higher frequencies; (b) The zoomed-in plot shows the modulation signal at 1*kHz* with an amplitude of 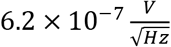, which corresponds to an external magnetic field of *H_m_* = 9.4 *μT*; (c) ME modulated voltage versus amplitude of the external magnetic field *H_m_* at 1*kHz* frequency, showing a LOD of 470pT; (d) LOD of ME antenna versus frequency of external alternating magnetic field *H_m_*. The LOD of the sensors improves with increasing frequency because, as it was shown in (a), 1/*f* electronics noise is dominant at lower frequencies. This noise decreases significantly with increasing frequency. At frequencies higher than 100*Hz*, both noise and ME modulation voltage decrease with almost the same rate, which is why the LOD, or i.e. the SNR, becomes almost flat. The inset plot shows the zoomed-in highlighted region (frequencies lower than 100Hz).

The diagram of experimental setup for magnetic sensing using an ME antenna is shown in Appendix E, Fig. S6. The ME antenna is excited using a lock-in amplifier through a directional coupler, and is operating with a 140 mV amplitude signal at its width resonance frequency of 63.6 MHz. The other port of the directional coupler, which is connected to the input of the lock-in amplifier for demodulation and post-processing, carries the reflected signal from ME antenna and contains the strain-mediated ME response of the FeGaB/AlN heterostructure. When excited (by RF voltage), the ME antenna is in presence of an external alternating magnetic field, and its reflected signal carries modulated components *ω_c_* – *ω_m_* and *ω_c_* + *ω_m_*, the amplitude of which is an indicator of the external magnetic field. A 37 Oe DC bias field provided by the larger Helmholtz coil (black color) is applied perpendicular to the length of the resonator in order to maximize the magnetoelectric coefficient and sensitivity. The reflected signal from the ME antenna is demodulated after passing through the input of the lock-in amplifier, and then its FFT is analyzed on a computer. Fig. 6a displays the power spectrum of the reflected signal from the ME antenna after demodulation. The plot shows both modulation components of *ω_c_* – *ω_m_* and *ω_c_* + *ω_m_* on two sides of the carrier signal that is located at the center and has an amplitude of 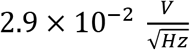. As it is shown, the 1/*f* electronics noise decreases significantly with increasing frequency, which improves the SNR of the sensor at higher frequencies. Fig. 6b shows the zoomed-in modulation signal of *ω_c_* + *ω_m_* at 1 kHz with an amplitude of 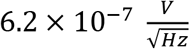 that corresponds to an external magnetic field of *H_m_* = 9.4 *μT*. Fig. 6c shows the ME modulated voltage versus amplitude of the external magnetic field *H_m_* at 1 *kHz* frequency. As discussed before in equation (4), the amplitude of the ME modulation signal changes linearly with external alternating magnetic field *H_m_*, making the device suitable to be used as a sensor. The plot shows that the limit of detection (LOD), or minimum detectable magnetic field, of the sensor is 470 pT. The ME modulation signal is at the same level of noise under magnetic fields smaller than LOD and therefore is lost in the noise. Fig. 6d shows the LOD of ME antenna versus frequency of external alternating magnetic field *H_m_*. As it was shown in Fig. 6a, the 1/*f* electronics noise of ME antenna decreases with increasing frequency; and the ME modulation voltage also decreases with frequency due to the fact that the coupling coefficient of ME heterostructure reduces with distance from the resonance frequency. At frequencies under 100 *Hz*, however, the rate of change of 1/*f* noise is faster than that of the ME coupling coefficient, which implies that the SNR of the ME antenna, and therefore the LOD, improves with increasing frequency as demonstrated in Fig. 6d. For modulation frequencies above 100 Hz on the other hand, both the 1/*f* noise and ME modulation signal decrease almost at the same rate, which is why the LOD of the sensor, i.e. the SNR, becomes almost flat. The inset plot in Fig. 6d shows the zoomed-in highlighted region (frequencies lower than 100 Hz), revealing a rapid change of the LOD at frequencies close to DC. Recent results in [23] show that the amplitude of the neural magnetic fields, for in-vivo measurement where the sensor is 10-100s of μm away from neural ensembles, is as high as 1-10 nT. In another work from Wikswo et al. [24], the authors reported a neural magnetic field with a frequency of approximately 1 kHz and an amplitude of 120 pT at 1.3 mm from the nerve. This value at this distance approximately agrees with the data in [23] where the modelling results in 126 pT at 1 mm, 1.3 nT at 100 μm, and 2.3 nT inside the neuronal ensemble. Therefore, the magnetic field measurements in these two works suggest that our ultra-compact smart ME antenna with a limit of detection of less than 470 pT has the capability to record neuronal magnetic fields.

## 4- Conclusion

We have reported a novel ultra-compact dual-band smart NEMS ME antenna that can be used for biomedical applications, in particular for implantable medical devices. The proposed dual-band ME antenna with a size of 250×174 μm^2^ has two acoustic resonance frequencies at 2.51 GHz and 63.6 MHz, where the former mode is used for wireless RF energy harvesting and the latter for low frequency magnetic field sensing, which can be used for neural recording. The wireless power transfer efficiency of the smart ME antenna is 1~2 orders of magnitude better than that of any other reported miniaturized micro-coils to date. The improved PTE allows the implantable device to operate at higher depth inside the body while adhering to the SAR limit set by the FCC. It was also shown that the ME antenna can be efficiently powered using an external Tx coil even if the device is misaligned or rotated after implantation. In addition, the smart ME antenna’s magnetic sensing mode shows an ultra-low limit of detection of less than 470 pT, which can be used for neuronal magnetic field sensing.

## Appendixes and Supplementary Materials

### A- Simulation Results of Strain and Displacement Distribution in FeGaB and AlN Thin-films

The results in Fig. S1 show the Multiphysics COMSOL simulation results (using coupled ACDC, Solid Mechanics, and Electrostatic modules) of strain and displacement distribution in FeGaB and AlN thin-films during energy harvesting process, where a 60nT magnetic field at 2.51GHz frequency is applied along the width of ME antenna. As the data in Fig. S1a and S1b show, this magnetic field creates a strain along the width and thickness directions of FeGaB material. Since the thickness of thin-films is significantly smaller compared to the width and length, the z-axis is scaled by x50 for a better visualization of strain distribution along the thickness. The induced strain in FeGaB layer is then transferred to the AlN thin-film, which is shown in Fig. S1c and S1d. The strain along the thickness mode of AlN induces a voltage at 2.51GHz frequency which can be used for energy harvesting purposes. Fig. S1e and S1f show the displacement along thickness direction of AlN thin-film at 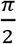 and 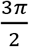 phases where displacement is maximum and minimum, respectively. The time domain animation of strain and displacement distribution over one full-cycle is also available in the supplementary materials as gif files.

**Fig. S1.**
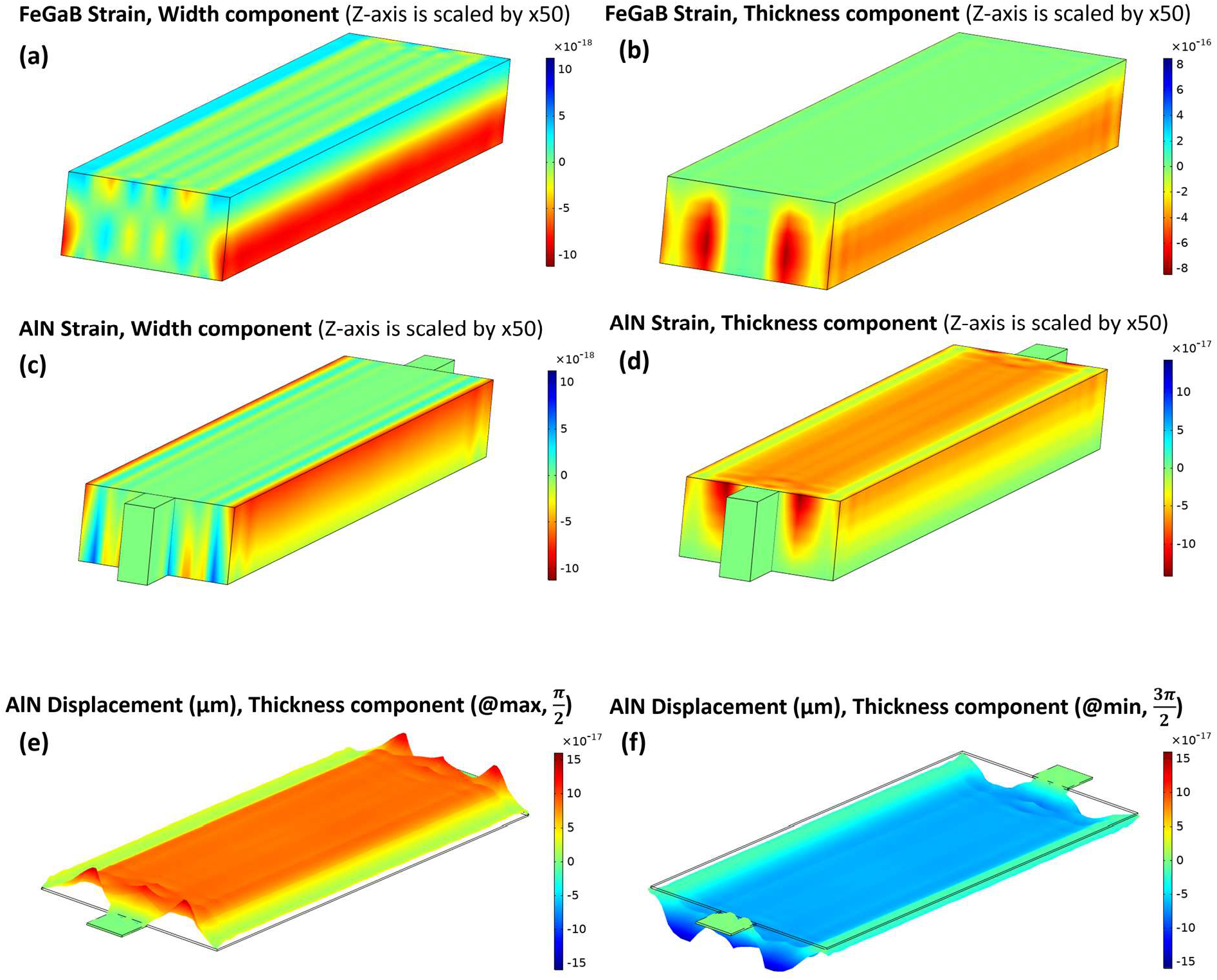
Strain and displacement distribution in FeGaB and AlN thin-films. (a) and (b) show the induced strain along the width and thickness of FeGaB layer, respectively; (c) and (d) show the induced strain along the width and thickness of AlN layer, respectively. The induced strain along the thickness mode of AlN thin-film generates a voltage at 2.51GHz frequency which can be used for energy harvesting applications. (e) and (f) show the displacement along thickness direction of AlN thin-film at 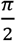 and 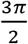 phase where displacement is maximum and minimum, respectively.

### B- Fabrication of a Smart NPR/FBAR ME Antenna

The fabrication process flow for the smart NPR/FBAR ME antenna is visualized in Fig. S2. The process starts with a high resistivity Silicon (Si) wafer (>10000 Ohm-cm). A 50 nm thick Pt film was sputter-deposited and patterned by lift-off on top of the Si substrate to define the bottom electrodes. Then, the 500 nm AlN film was sputter-deposited and vias were etched with H_3_PO_4_ to access the bottom electrodes. Afterwards, the AlN film was etched by Inductively Coupled Plasma (ICP) etching using Cl_2_ based chemistry to define the shape of the resonant nano-plate. Next, a 100 nm thick gold (Au) film was evaporated and patterned by lift-off to form the top ground. Finally, 500 nm thick FeGaB/Al_2_O_3_ multilayer layer was deposited by a magnetron sputtering and patterned by lift-off. A 100 Oe in-situ magnetic field bias was applied during the magnetic deposition along the width direction of the device to pre-orient the magnetic domains. Then, the structure was released by XeF2 isotropic etching of the Silicon substrate. Fig. S2f shows the device layout of the ME FBAR antenna with the detailed dimensions.

The magnetic multilayer with the structure of [FeGaB (45 nm)/Al_2_O_3_ (5 nm)] ×10 was sputter-deposited on AlN thin film with a Ta (5nm) seed layer at the Ar atmosphere of 3 mTorr with a background pressure of <1×10^−7^ Torr. The Ta seed layer promoted the FeGaB thin film growth exhibiting narrow resonance linewidth and close-to-bulk magnetic moment. The FeGaB layer was co-sputtered from FeGa (DC sputtering) and B (RF sputtering) targets. The Al_2_O_3_ layer was deposited by RF sputtering using an Al_2_O_3_ target. The deposition rates are calibrated by X-ray reflectivity.

**Fig. S2.**
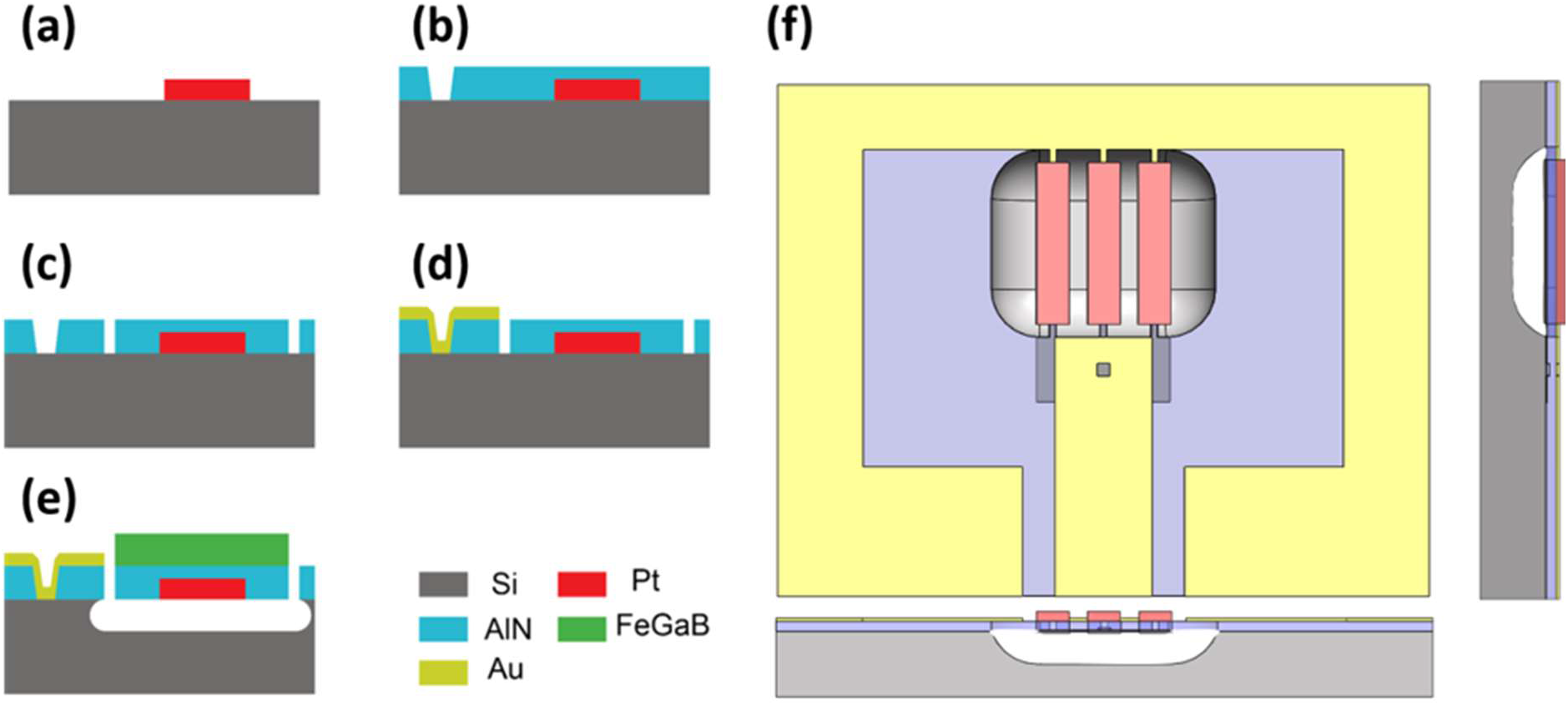
Smart NPR/FBAR ME antenna fabrication process: (a) Pt thin film deposition and patterning. (b) AlN thin film deposition and wet etch of vias. (c) AlN thin film ICP etching. (d) Top electrode Au deposition and patterning. (e) Magnetic multilayer FeGaB/AlO_x_ deposition and pattering followed by Si substrate release. (f) Device layout of the ME FBAR antenna.

The reason we have used FeGaB/Al2O3 multilayers (discussed in Appendix B, fabrication of ME antenna) is that they demonstrate eddy-current loss reduction, lower out of plane anisotropy, and enhance permeability in comparison with a single FeGaB layer with the same thickness. Therefore, we are not worried about the eddy-current loss. We have been extensively researching ME antennas and ME sensors optimization since 2015, and we are one of the leading teams on this effort. As for the electrode, we have tried three different approaches: (1) Using FeGaB film as an electrode; (2) Using gold between FeGaB and AlN for better conductivity; (3) Using gold as an electrode on top of the FeGaB for better conductivity. The results always showed better performance, sensitivity, and quality factors using FeGaB directly as the electrode. Therefore, in this project, we used FeGaB directly instead of using other metal on top or in-between.

### C- Misalignment and Rotation of the ME Antenna in other Directions (for Energy Harvesting Tests)

**Fig. S3.**
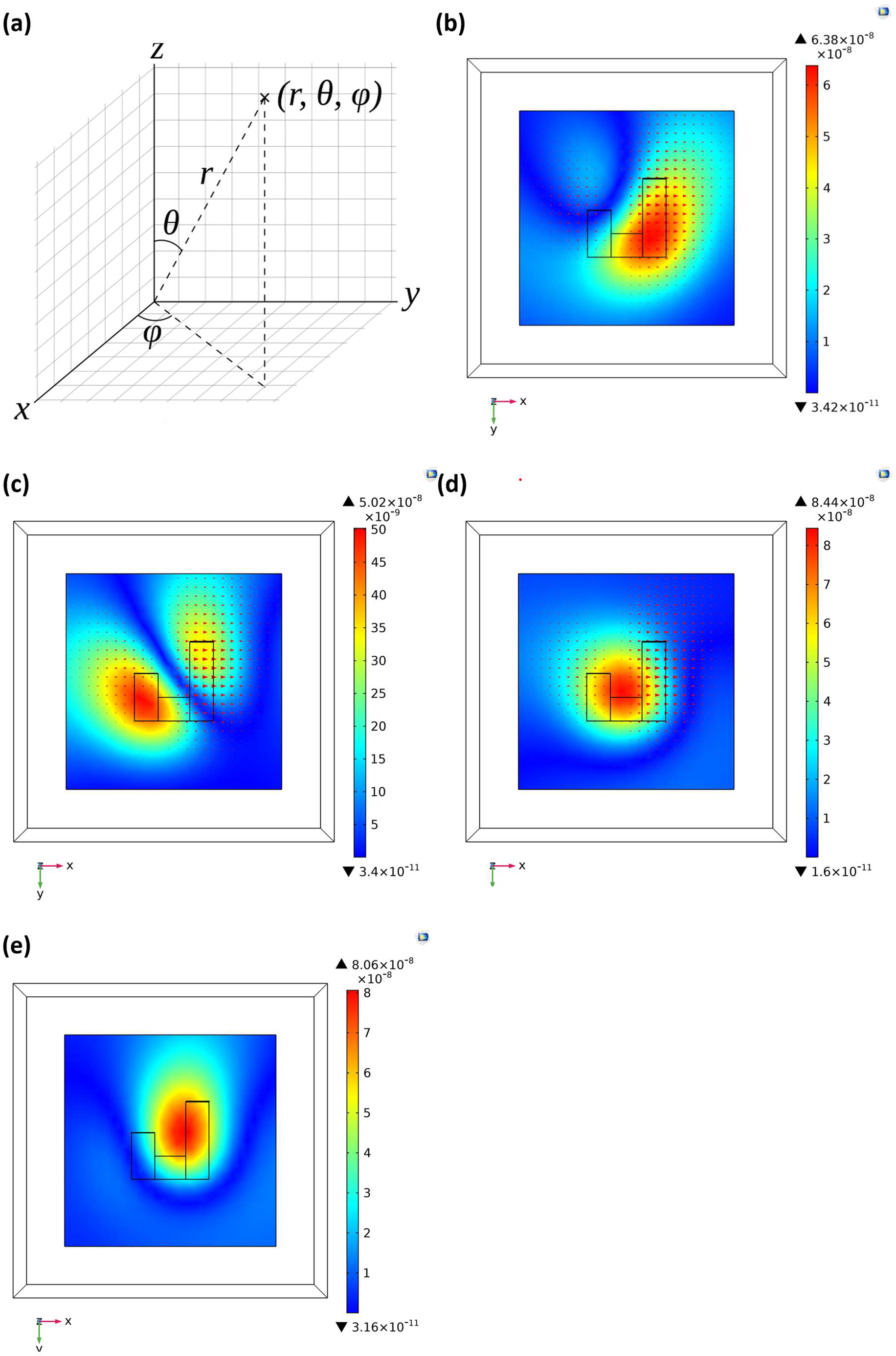
(a) The spherical coordinate systems used for demonstrating different orientations, where the vector 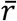 shows the direction towards which the ME antenna is sensitive. *ϕ* and *θ* show the angles between 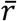 vector and *x* and *z* axis, respectively; (b) the orientation in which *ϕ* = 45° and *θ* = 90°; (c) the orientation in which *ϕ* = 135° and *θ* = 90°; (d) the orientation in which *ϕ* = 90° and *θ* = 45°; (e) the orientation in which *ϕ* = 0 and *θ* = 45°.

The magnetic flux density components (Bx, By, Bz) generated by Tx coil in the XY plane were shown and discussed in Section 2. In this appendix, we will discuss the magnetic flux density distribution in several other orientations. Fig. S3a shows the spherical coordinate system used for demonstrating different orientations, where the vector 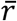 shows the direction towards which ME antenna is sensitive. *ϕ* and *θ* are the angles between 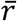 vector and the *x* and *z* axes, respectively. Fig. S3b shows the orientation in which *ϕ* = 45° and *θ* = 90°, i.e. this is the case where the ME antenna is 45° between the x- and y-axis. This orientation was discussed with more in details in Section 2. Fig. S3c shows the orientation in which *ϕ* = 135° and *θ* = 90°. Fig. S3d shows the orientation in which *ϕ* = 90° and *θ* = 45°. Fig. S3e shows the orientation in which *ϕ* = 0 and *θ* = 45°.

### D- Magnetic Field and Eddy Current Distribution Inside the Tissue and Air Mediums

As discussed in Section 2 of the article, the magnetic field distribution is different in air and tissue mediums. According to simulation results, eddy current loops generated inside the tissue impact the magnetic field distribution. These eddy current loops are generated by the initial magnetic flux density generated in the Tx coil, which are flowing towards the z-axis. Fig. S4a and S4b show the magnetic flux density on two different planes of air and tissue mediums, respectively. The vertical planes are the same cross-section planes shown in Fig. S4c and S4d. They show that the magnetic field distribution is normal in air but distorted in tissue medium. There are two vortexes in tissue medium, which are due to eddy current loops created in the meninges layer because of its higher conductivity. The horizontal planes show the magnetic flux density above the coil at the interface of meninges and grey matter layers, where the mentioned vortexes are created. Fig. S4c and S4d, horizontal planes, show the current density (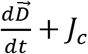, the first and second terms are displacement current density and conduction current density, respectively) on the interface of meninges and grey matter layers in air and tissue medium, respectively. The magnetic flux Bz, shown in Fig. S4b, creates eddy current loops rotating around the B-fields in the tissue. These eddy current loops in turn generate the magnetic field loops on the vertical plane which lead to the vortexes shown with black-loops. There are no generated eddy current loops in the air, and therefore the Tx coil field distribution is normal and not distorted.

**Fig. S4.**
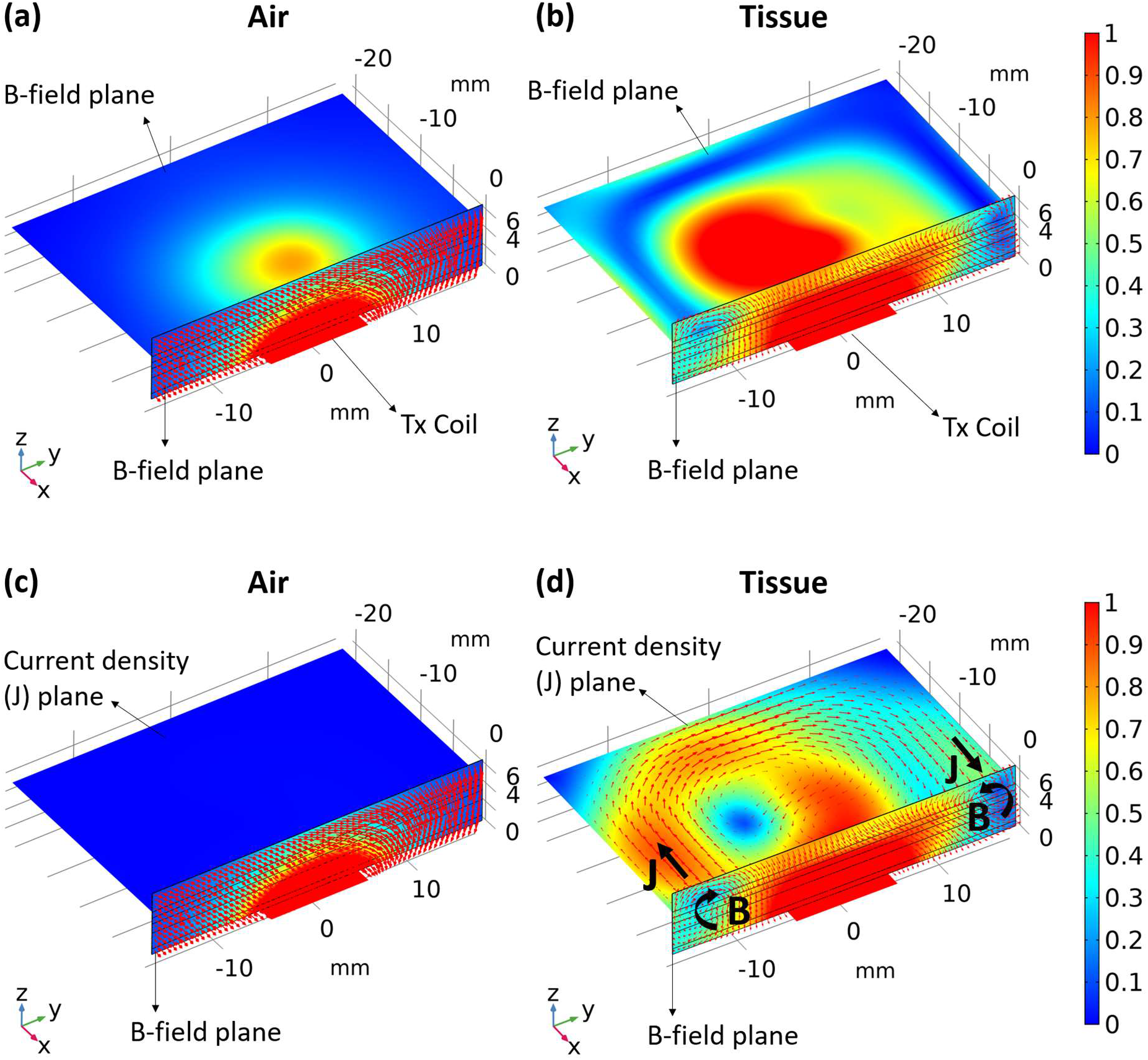
(a) and (b) magnetic flux distribution on two vertical and horizontal cross-section planes in air and tissue mediums, respectively. The fields on horizontal planes are mostly towards z-axis; (c) and (d) show the current density on horizontal plane and magnetic flux density on vertical plane in the air and tissue mediums, respectively. The horizontal plane is the interface of meninges and grey matter layers. As it is shown the magnetic field Bz create eddy current loops on the horizontal plane in fig S3d. These eddy currents in turn generate magnetic field loops which interfere with the magnetic fields from Tx coil. Therefore, they distort the initial field distribution and create the two vortexes shown in the vertical plane.

### E- Energy Harvesting and Magnetic Field Sensing Experimental Setup

The experimental setup for the energy harvesting measurements is displayed in Fig. S5. The Tx coil and ME antenna are mounted on two 3D printed plastic manipulators in order to precisely adjust their position. Plastic manipulator, instead of metallic ones, are used to minimize the effect of objects surrounding the Tx coil and their impact on S_11_ of the coil.

The diagram of experimental setup for magnetic sensing using ME antenna is shown in Fig. S6. The zoomed-in part on the right shows the device under test and its orientation with respect to the external magnetic field. The part number of the coupler used in the experiment is ZDC-10-1+ from Mini-Circuits. A 37 Oe DC bias field provided by the larger Helmholtz coil (black color) is applied perpendicular to the length of the resonator in order to maximize the magnetoelectric coefficient and sensitivity. The alternating magnetic field *H_m_*, which is the external test magnetic field, is in the same direction as DC bias field and is provided by smaller Helmholtz coil (blue color). The larger and smaller Helmholtz coils are driven by a Keithley 2461 DC current source and Keithley 6221 AC current source, respectively.

**Figure S5.**
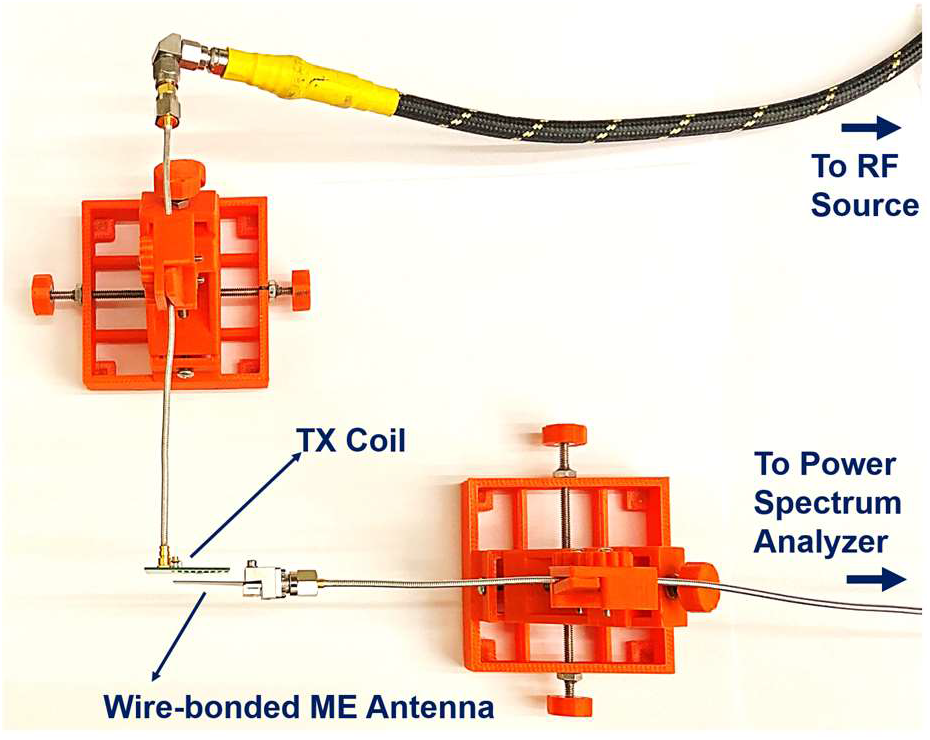
Experimental setup used for the energy harvesting measurements.

**Figure S6.**
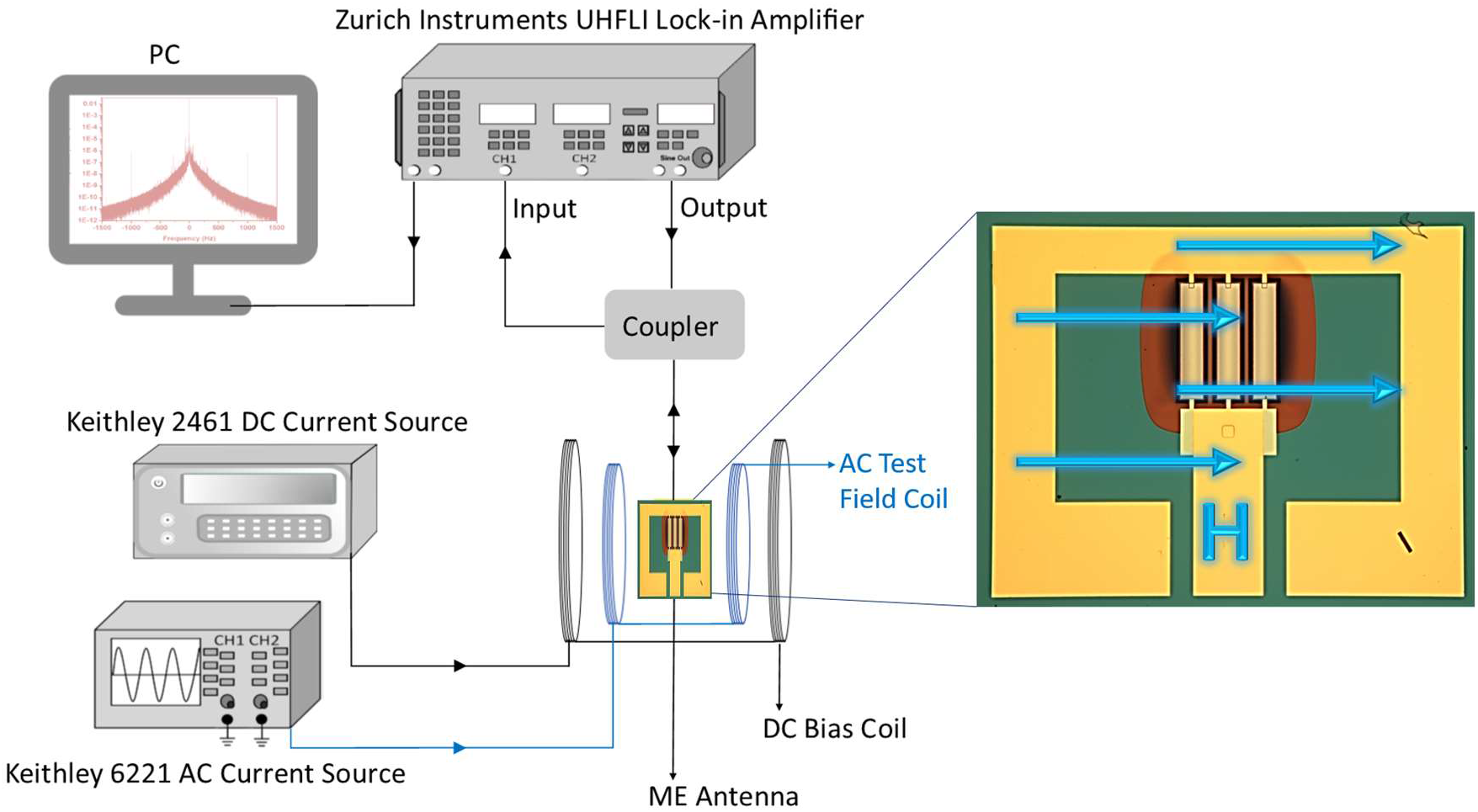
Experimental setup used for magnetic sensing using an ME antenna, where the zoomed-in part on the right shows the device under test and its orientation with respect to external magnetic field. It is notable that this is the same direction as during the energy harvesting tests.

